# Copy number variation in fungi and its implications for wine yeast genetic diversity and adaptation

**DOI:** 10.1101/233122

**Authors:** Jacob L. Steenwyk, Antonis Rokas

**Affiliations:** Department of Biological Sciences, Vanderbilt University, Nashville, TN 37235, USA

**Keywords:** Structural variation, alcohol fermentation, sugar metabolism, gene duplication, gene loss, population genomics

## Abstract

In recent years, copy number (CN) variation has emerged as a new and significant source of genetic polymorphisms contributing to the phenotypic diversity of populations. CN variants are defined as genetic loci that, due to duplication and deletion, vary in their number of copies across individuals in a population. CN variants range in size from 50 base pairs to whole chromosomes, can influence gene activity, and are associated with a wide range of phenotypes in diverse organisms, including the budding yeast *Saccharomyces cerevisiae.* In this review, we introduce CN variation, discuss the genetic and molecular mechanisms implicated in its generation, how they can contribute to genetic and phenotypic diversity in fungal populations, and consider how CN variants may influence wine yeast adaptation in fermentation-related processes. In particular, we focus on reviewing recent work investigating the contribution of changes in CN of fermentation-related genes associated with the adaptation and domestication of yeast wine strains and offer notable illustrations of such changes, including the high levels of CN variation among the *CUP* genes, which confer resistance to copper, and the preferential deletion and duplication of the *MAL1* and *MAL3* loci, respectively, which are responsible for metabolizing maltose and sucrose. Based on the available data, we propose that CN variation is a substantial dimension of yeast genetic diversity that occurs largely independent of single nucleotide polymorphisms. As such, CN variation harbors considerable potential for understanding and manipulating yeast strains in the wine fermentation environment and beyond.

## Introduction

Genetic variation in natural populations is shaped by diverse biological processes, such as genetic drift and natural selection (Chakravarti, 1999), and is, in part, responsible for phenotypic variation. For example, arginine auxotrophy in the baker’s yeast *Saccharomyces cerevisiae* is a Mendelian inherited trait due to polymorphisms in the *ARG4* locus (Brauer et al., 2006), whereas variation in *S. cerevisiae* colony morphology is a complex trait driven by variants in several different genes (Taylor et al., 2016). The aforementioned yeast phenotypes are all caused by SNPs or small insertions and deletions, which are by far the most well characterized types of genetic variation not only in yeast, but in any kind of organism (McNally et al., 2009; Sachidanandam et al., 2001; Schacherer et al., 2009). In recent years, however, several studies in diverse organisms have revealed that genomes also harbor an abundance of structural variation, which too contributes to populations’ genetic and phenotypic diversity (Stranger et al., 2007; Zhang et al., 2009).

Variation in the structure of chromosomes, or structural variation, encompasses a wide array of mutations including insertions, inversions, translocations, and copy number (CN) variants (i.e., duplications and deletions) (Feuk et al., 2006) and, in humans, accounts for an estimated average of 74% of the nucleotide differences between two genomes (Rahim et al., 2008). The major influence of several types of structural variation, such as large-scale inversions, translocations, and insertions, on phenotype is better understood because many such variants can be microscopically examined and lead to classic human genetic disorders, such as Down’s syndrome (Gu et al., 2016; Rausch et al., 2012; Youings et al., 2004). In contrast, many CN variants are submicroscopic and eschewed attention until the advent of whole genome sequencing technologies (Feuk et al., 2006).

CN variants are defined as duplications or deletions that range from 50 base pairs to whole chromosomes (Figure 1) and can significantly influence phenotypic diversity (Arlt et al., 2014; Zhang et al., 2009). For example, in humans, the CN of the salivary amylase gene, *AMY1,* is higher in populations with high-starch diets and correlated with salivary protein abundance thereby improving digestion of starchy foods (Perry et al., 2007). Levels of CN variation have been examined in diverse organisms across the tree of life, including animals (e.g., Humans; *Homo sapiens*: Sudmant et al., 2015, House mouse; *Mus musculus*: Pezer et al., 2015), plants (e.g., soybean; *Glycine max:* Cook et al., 2012, maize; *Zea mays:* Swanson-Wagner et al., 2010) and fungi (e.g., *Cryptococcus neoformans:* Hu et al., 2011, *Batrachochytrium dendrobatidis:* Farrer et al., 2013, *Zymoseptoria tritici:* Hartmann and Croll, 2017).

**Figure 1.**
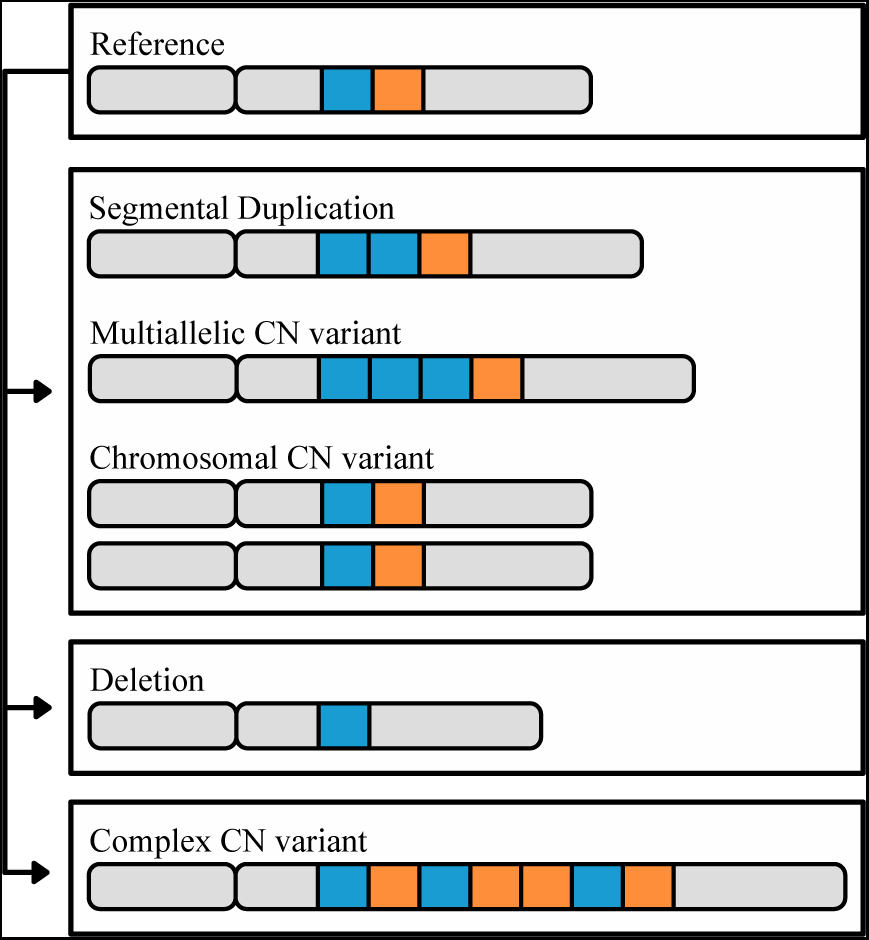
The different types of CN variation. CN variants range in size (50 base pairs or greater) to whole chromosomes, and are identified through comparison to a reference genome. In this cartoon, a reference chromosome containing two highlighted loci, in blue and orange, is shown on top. The second chromosome illustrates an example of a segmental duplication CN, in which there are two copies of the blue locus. The third chromosome illustrates an example of a multiallelic CN variant, where the duplicated locus contains 3 or more copies. The fourth pair of chromosomes illustrates a CN variant associated with the duplication of an entire chromosome. Finally, the last two chromosomes illustrate deletion and complex CN variants, respectively; deletion CN variants are associated with loci that are not present relative to the reference, and complex CN variants refer to a combination of duplications, deletions, insertions, and/or inversions relative to the reference.

*S. cerevisiae* has been an important model for genetics, genomics and evolution (Botstein et al., 1997; Goffeau et al., 1996; Winzeler et al., 1999). Much of what we know about the evolutionary history of *S. cerevisiae* stems from investigating genome-wide patterns of SNPs among globally distributed strains. Examination of genome-wide patterns of SNP variation has yielded valuable insights into yeast function in the wine fermentation environment. For example, 13 SNPs in *ABZ1,* a gene associated with nitrogen biosynthetic pathways, have been shown to modify the rate of fermentation and nitrogen utilization during fermentation (Ambroset et al., 2011).

Interrogations of genome-wide patterns of SNPs have also shown that industrial clades - including those of beer, bread, cacao, sake, and wine - often mirror human history (Cromie et al., 2013; Gallone et al., 2016; Gonçalves et al., 2016; Schacherer et al., 2009; Sicard and Legras, 2011), suggesting that human activity has greatly influenced *S. cerevisiae* genome evolution (Yue et al., 2017). Furthermore, SNP-based studies have repeatedly found that wine strains of *S. cerevisiae* exhibit low levels of genetic diversity (Borneman et al., 2016; Cromie et al., 2013; Liti et al., 2009; Schacherer et al., 2009; Sicard and Legras, 2011), consistent with a historical population bottle-neck event that reduced wine yeast genetic variation. The low SNP diversity among wine yeast strains has led some to suggest that wine strain development may benefit from the introduction of genetic variation from yeasts outside the wine clade (Borneman et al., 2016). However, recent studies examining CN variation among wine associated strains of *S. cerevisiae* have identified considerable genetic diversity (Gallone et al., 2016; Gonçalves et al., 2016; Steenwyk and Rokas, 2017), suggesting that standing CN variation in wine strains may be industrially relevant.

In the present review, we begin by surveying the molecular mechanisms that lead to CN variant formation, we next discuss the contribution of CN variation to the genetic and phenotypic diversity in fungal populations, and close by examining the CN variation in wine yeasts and the likely phenotypic impact of CN variants in the wine fermentation environment.

## Copy number variation and the molecular mechanisms that generate it

Copy number (CN) variants, a class of structural variants, are duplicated or deleted loci that range from 50 base pairs (bp) to whole chromosomes in length (Figure 1) and have a mutation rate 100-1,000 times greater than SNPs (Arlt et al., 2014; Sener, 2014; Zhang et al., 2009). CN variable loci can in turn be broken down into three subclasses (Figure 1) (Estivill and Armengol, 2007). The first subclass encompasses variants that originate via duplications; in the genome, these can appear as either identical or nearly identical copies, or multi-allelic CN variants (Bailey and Eichler, 2006; Usher and McCarroll, 2015). The extreme version of this subclass are chromosomal CN variants that correspond to duplications of entire chromosomes. The second subclass encompasses CN variants that originate via deletion leading to the loss of the sequence of a locus in the genome. The third subclass includes complex CN variants where a locus exhibits a combination of duplication, deletion, insertion, and inversion events (Usher and McCarroll, 2015).

CN variants are commonly generated from aberrant DNA repair via three mechanisms: homologous recombination (HR), non-homologous repair (NHR), and environmental stimulation (Figure 2) (Hastings et al., 2009b; Hull et al., 2017). HR is a universal process associated with DNA repair and requires high sequence similarity across 60 - 300 bps (Hua et al., 1997; Petukhova et al., 1998). HR is initiated by double-strand breaks caused by ionizing radiation, reactive oxygen species, and mechanical stress on chromosomes such as those associated with collapsed or broken replication forks (Aylon and Kupiec, 2004; Hastings et al., 2009b; Khanna and Jackson, 2001). Improper repair by HR can result in duplication, deletion, or inversion of genetic material (Reams and Roth, 2015). Non-allelic HR (also known as ectopic recombination), defined as recombination between two different loci of the same or different chromosomes that share sequence similarity and are ≥300 base pairs in length, is among the most well-studied examples of improper repair (Kupiec and Petes, 1988; Prado et al., 2003). Most evidence of non-allelic HR resulting in CN variation is directly associated with low copy repeats or transposable elements (Hurles, 2005; Xu and Boeke, 1987). For example, a duplication and deletion may result during unequal crossing over of homologous sequences (Figure 2a) (Carvalho and Lupski, 2016). Improper HR may also occur at collapsed or broken replication forks by break-induced replication (BIR) (Figure 2b). BIR requires 3’ strand invasion at the allelic site of stalled replication to properly restart DNA synthesis (Figure 2bi) (Llorente et al., 2008), however, template switching, the non-allelic pairing of homologous sequences, in the backward (Figure 2bii) or forward (Figure 2biii) direction can result in a duplication or deletion, respectively (Morrow et al., 1997; Smith et al., 2007). Although HR occurs with high fidelity, errors in the process, which are thought to increase in frequency during mitosis and meiosis, can generate CN variants (Hastings et al., 2009b).

**Figure 2.**
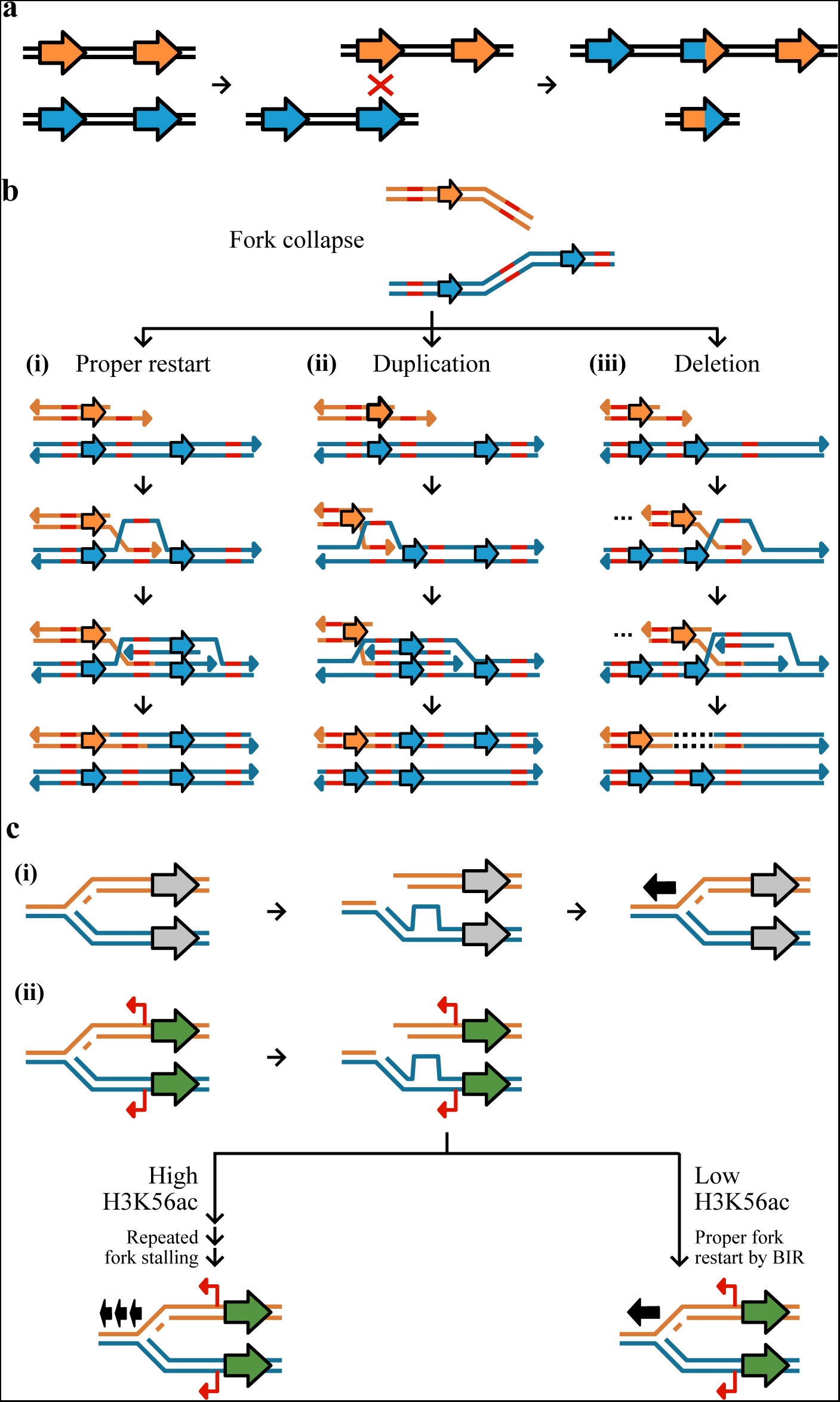
Mechanisms of CN variant formation. CN variants typically occur as a result of aberrant replication via homologous recombination, non-homology based mechanisms, and environmentally stimulated processes. (a) Unequal crossing over during recombination may result in duplication and deletion. Here, two equal strands of DNA with two genes (represented by the orange or blue arrows) have undergone unequal crossing over due to the misalignment of a homologous sequence. This results in one DNA strand having three genes and the other one gene. (b and c) A major driver of CN variant formation is aberrant DNA replication. (b, top) Double strand breaks at replication forks or collapsed forks are often repaired via Break-induced replication (BIR). (bi) Proper BIR starts with strand invasion of a homologous or microhomologous sequence (shown in red) to allow for proper fork restart. (bii) If template switching occurs in the backward direction, a segment of DNA will have been replicated twice resulting in a duplication; (biii) in contrast, template switching in the forward direction results in a deletion represented by a dashed line in the DNA sequence. Erroneous BIR may be mediated by microhomologies as well. (c) CN variants may be stimulated near genes that are highly expressed due to an increased chance of fork stalling. (ci) If a replication fork breaks down near a gene that is not expressed (grey) and restarts once (represented by one black arrow), no mutation will occur. (cii) If a replication fork breaks down near a gene that is expressed (green) with cryptic unstable transcripts (red) then there may be two outcomes dependent on the degree of the H3K56ac acetylation mark. If there are low levels of H3K56ac, it is more likely that there will be proper fork restart by BIR (represented by one black arrow). If there are high levels of H3K56ac, it is more likely that there will be repeated fork stalling (represented by three black arrows) (see figure 8 from Hull *et al.* 2017).

In contrast to HR, NHR utilizes microhomologies (typically defined as ~65% or more sequence similarity of short sequences up to ten bases long) or does not require homology altogether, and can too lead to CN variant formation (Daley et al., 2005; McVey and Lee, 2008). NHR can occur by two mechanisms: non-replicative and replicative (Hastings et al., 2009b). Non-replicative mechanisms include non-homologous end joining and microhomology-mediated end-joining (Lieber, 2008; McVey and Lee, 2008). Non-homologous end-joining refers to the direct ligation of sequences in a double-strand break (Daley et al., 2005). Prior to ligation, there may be a loss of genetic material or the addition of free DNA (e.g., from transposable elements or mitochondrial DNA) (Yu and Gabriel, 2003). Microhomology-mediated end joining is similar to non-homologous end-joining but occurs more frequently, requires different enzymes, and leverages homologies 1-10 base pairs in length to ensure more efficient annealing (Lieber, 2008; Yu et al., 2004). Non-homologous end-joining and microhomology-mediated non-homologous end-joining are primarily associated with small insertions and deletions and therefore are not likely to be a major driver of CN variation (Gu et al., 2008; Yu and Gabriel, 2003). Replicative mechanisms of CN variant formation include replication slippage, fork stalling, and microhomology BIR. Replication slippage occurs along repetitive stretches of DNA resulting in the duplication or deletion of sequence between repetitive regions (Hastings et al., 2009b). Fork stalling is thought to cause large CNVs of 20 kb average length through template switching between distal replication forks rather than within a replication fork (Slack et al., 2006).

However, fork stalling without distal template switching can also be highly mutagenic and induce CN variants (Hull et al., 2017; Paul et al., 2013). Lastly, microhomology-mediated break-induced replication occurs when the 3’ end of a collapsed fork anneals with any single-stranded template that it shares microhomology with to reinitiate DNA synthesis (Figure 2b) (Hastings et al., 2009b). Annealing can occur in the backward (Figure 2bii) or forward (Figure 2biii) direction of the allelic site causing a duplication or deletion, respectively, and is thought to be the primary cause of low copy repeats (Hastings et al., 2009a).

The third mechanism is associated with an epigenetic mark that can stimulate the formation of CN variants. Histone acetylation, specifically H3K56ac, is, in part, environmentally driven (Turner, 2009), associated with highly transcribed loci, and can promote CN variant formation through repeated fork stalling or template switching (Figure 2c) (Hull et al., 2017). For example, it has been shown that exposure to environmental copper stimulates the generation of CN variation in *CUP1,* a gene that is associated with copper resistance when duplicated (Fogel and Welch, 1982), thereby increasing the likelihood of favorable alleles that exhibit increased copper resistance (Hull et al., 2017). Similarly, environmental formaldehyde exposure was shown to stimulate CN variation (Hull et al., 2017) of the *SFA1* gene, which confers formaldehyde resistance at higher CNs (Wehner et al., 1993). Altogether, these experiments provide insight to how perturbations of an environmental parameter may stimulate CN variation at a locus important to adaptation in the new environment (Hull et al., 2017).

## Copy number variation as a source of phenotypic diversity

CN variants can have multiple effects on gene activity, such as changing gene dosage (i.e., gene CN; Figure 3) and interrupting coding sequences (Itsara et al., 2009; Sener, 2014). These effects can be substantial; for example, 17.7% of gene expression variation in human populations can be attributed to CN variants (Stranger et al., 2007). Furthermore, changes in human gene expression attributed to CN variants have little overlap with changes in gene expression caused by SNPs, suggesting the two types of variation independently affect gene expression (Stranger et al., 2007). Additionally, gene CN tends to correlate with levels of both gene expression and protein abundance (Henrichsen et al., 2009; Perry et al., 2007; Stranger et al., 2007). For example, changes in gene expression and therefore protein abundance caused by chromosomal CN variation in human chromosome 21 are thought to contribute to Down syndrome (Aivazidis et al., 2017; Kahlem et al., 2004).

**Figure 3.**
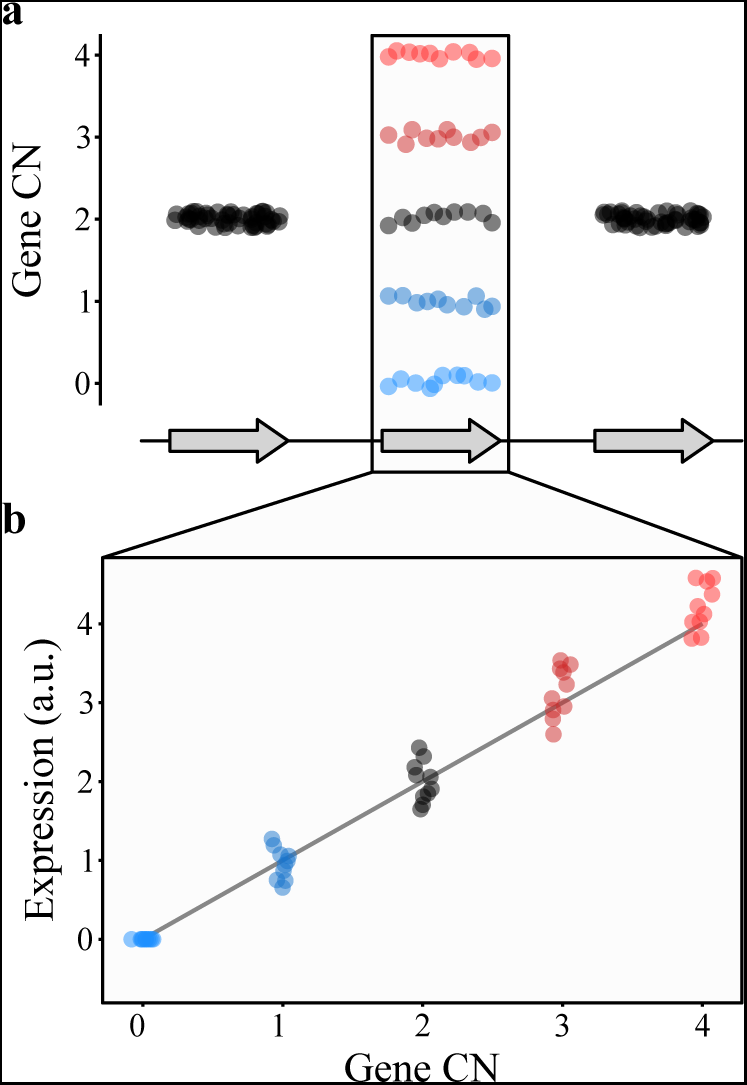
CN variation can alter gene expression. (a) Consider a gene whose CN ranges from 0 to 4 (blue to black to red) among individuals (represented by dots) in a population (middle gene). (b) Generally, CN and gene expression (represented as arbitrary units or a.u.) correlate with one another such that individuals with lower CN values will have lower levels of gene expression of that gene while those with higher CN values will have higher levels of gene expression.

## Copy number variation as a source of genetic and phenotypic diversity in fungal populations

CN variant loci contribute to population genetic and phenotypic diversity (Box 1), such as virulence (Farrer et al., 2013; Hu et al., 2011b), in diverse fungal species, including as the baker’s yeast *Saccharomyces cerevisiae* (ASCOMYCOTA, Saccharomycetes) (Gallone et al., 2016; Gonçalves et al., 2016; Steenwyk and Rokas, 2017), the fission yeast *Schizosaccharomycespombe* (ASCOMYCOTA, Schizosaccharomycetes) (Jeffares et al., 2017), the human fungal pathogen *Cryptococcus deuterogattii* (BASIDIOMYCOTA, Tremellomycetes) (previously known as *Cryptococcus gattii* VGII; Steenwyk et al., 2016) and *C. neoformans* (Hu et al., 2011b), the amphibian pathogen *Batrachochytrium dendrobatidis* (CHYTRIDIOMYCOTA, Chytridiomycetes) (Farrer et al., 2013), and the wheat pathogen *Zymoseptoria tritici* (ASCOMYCOTA, Dothideomycetes) (Hartmann and Croll, 2017).

#### Box 1.

Standard population genetic principles of shifts in allele frequencies (Felsenstein, 1976; Moritz, 1994) can be applied to CN variants. To illustrate the case, we provide an example using the *CUP1* locus, where high CN provides protection against copper poisoning (Fogel and Welch, 1982), of how the allele frequency of a CN variant can increase through its phenotypic effect. Suppose that in a yeast population exposed to copper that all individuals do not harbor CN variation at the *CUP1* locus. Through a mutational event, a beneficial *CUP1* allele that contains two or more copies of the locus may appear in the population. (a) Yeast with two or more copies of *CUP1,* which in turn lead to higher *CUP1* protein levels, will be better and more efficient at copper sequesteration unlike the parental allele and therefore avoiding copper poisoning (Fogel and Welch, 1982). (b) Assuming a large population size and strong positive selection, changes in allele frequency will occur in the population due to changes in yeast survivability and ability to propagate. More specifically, the frequency of the beneficial allele (i.e., *CUP1* duplications) will increase depending on the strength of selection, which increases as the concentration of environmental copper increases, and the parental allele will decrease.

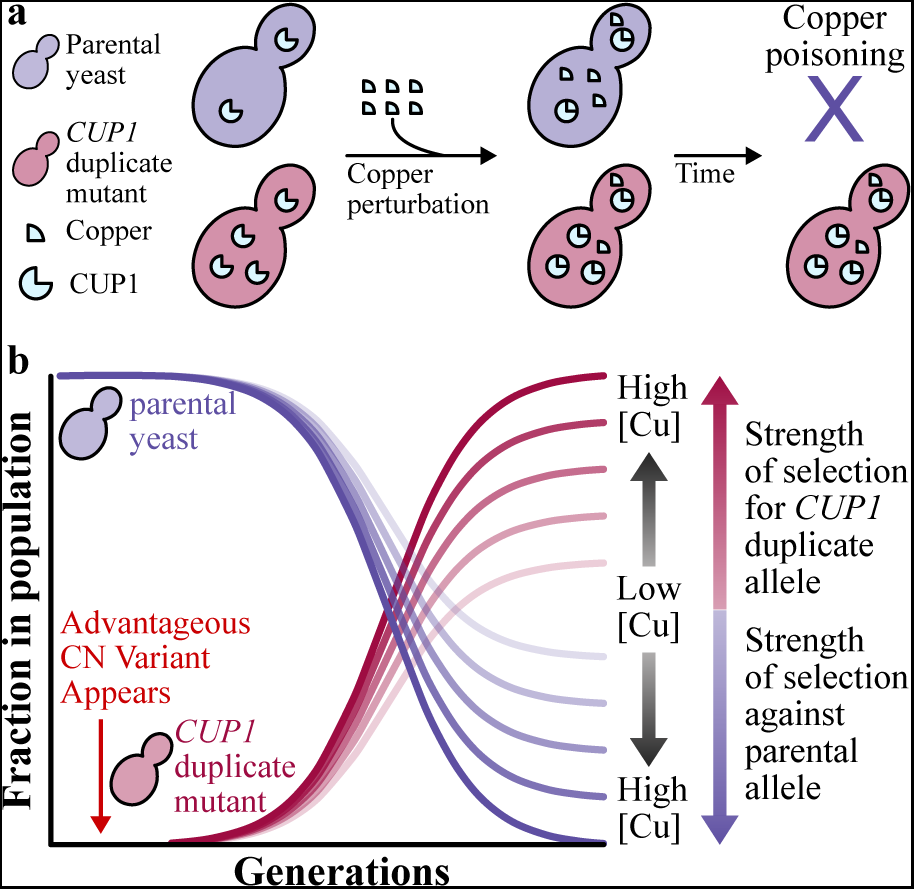

Importantly, the degree of CN variation (which can be represented by CN variable base pairs per kilobase) in fungal populations is not always correlated to the degree of SNP variation (which can be represented by SNPs per kilobase) (Figure 4a). For example, there is no correlation between CN variable base pairs per kilobase and SNPs per kilobase among *S. cerevisiae* wine strains (Steenwyk and Rokas, 2017) and a population of *Cryptococcus deuterogattii* (Steenwyk et al., 2016). Interestingly, both populations harbor low levels of SNP diversity; for *S. cerevisiae* wine strains this is due to a single domestication-associated bottleneck event (Cromie et al., 2013; Liti et al., 2009; Schacherer et al., 2009; Sicard and Legras, 2011), whereas for *C. deuterogattii* this is because the samples stem from three clonally evolved subpopulations from the Pacific Northwest, United States (Engelthaler et al., 2014). In contrast, a significant correlation is observed between CN variable base pairs per kilobase and SNPs per kilobase among individuals in a globally distributed population of *S. pombe* (Jeffares et al., 2015).

**Figure 4.**
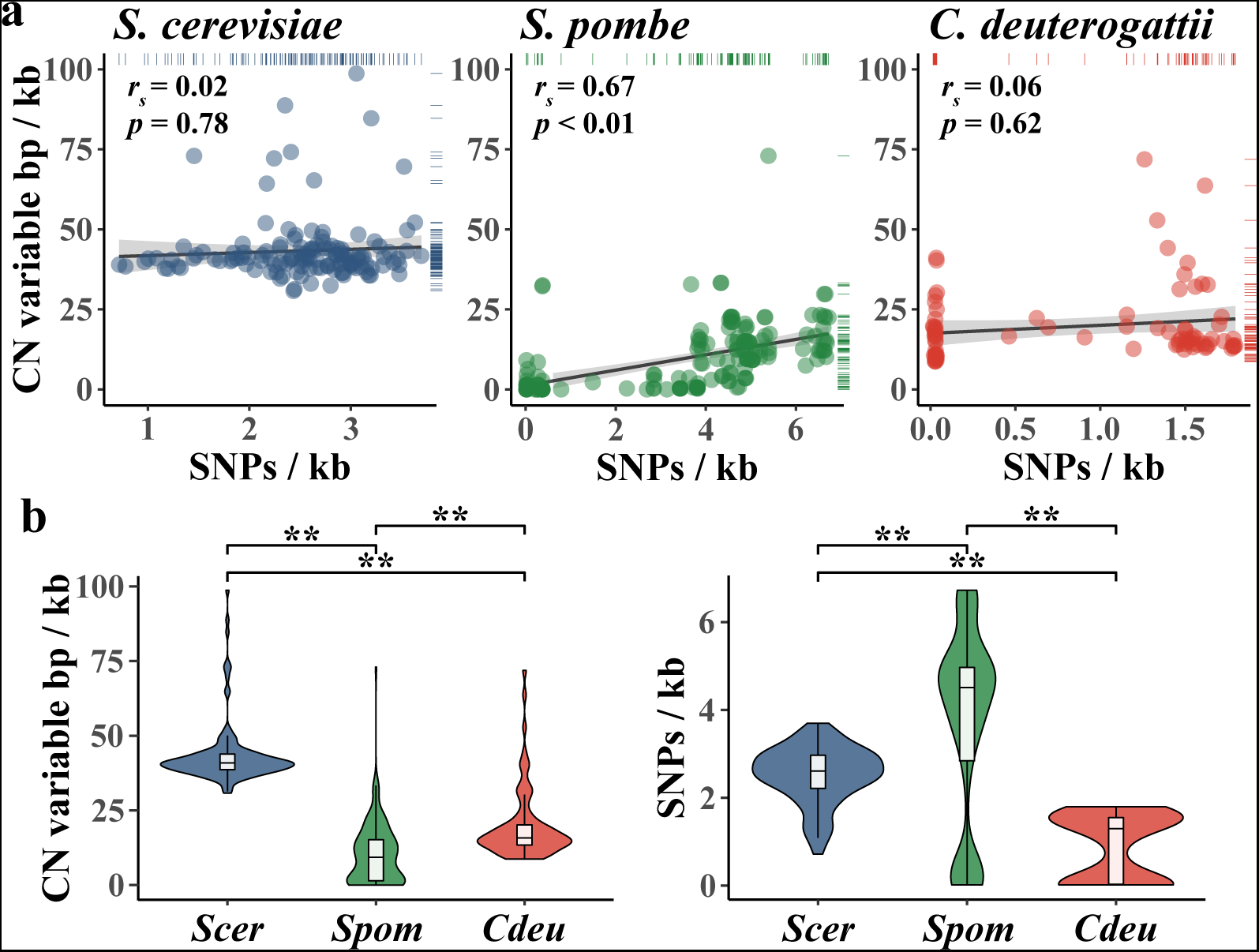
Comparison of genomic content affected by CN variants and SNPs in 3 fungal species. (A) SNPs per kb is not significantly correlated with CN variable base pairs per kb in *S. cerevisiae* wine strains (blue; *r_s_* = 0.02; *p* = 0.78; Spearman rank correlation) and *C. deuterogattii* (red; *r_s_* = 0.06; *p* = 0.62; Spearman rank correlation); the reverse is true in *S. pombe* (green; *r_s_* = 0.67; *p* < 0.01; Spearman rank correlation). (b, left) CN variable base pairs per kb in wine strains of *S. cerevisiae* is greater than *C. deuterogattii* and *S. pombe* (*p* < 0.01; Kruskal-Wallis and *p* < 0.01 for all Dunn’s test pairwise comparisons with Benjamini-Hochberg multi-test correction). (b, right) SNPs per kb is low among *S. cerevisiae* wine strains (*Scer*) compared to *S. pombe* (*Spom*) but greater than a clonally expanded population of *C. deuterogattii* (*Cdeu*) (*p* < 0.01; Kruskal-Wallis and *p* < 0.01 for all Dunn’s test pairwise comparisons with Benjamini-Hochberg multi-test correction). Data from Jeffares et al., 2015, 2017 (*Spom*); Steenwyk et al., 2016 (*Cdeu*); Steenwyk and Rokas, 2017 (Scer).

The proportion of the genome exhibiting CN and SNP variation also varies across *S. cerevisiae, S. pombe,* and *C. deuterogattii* populations. For example, CN variable base pairs per kilobase are significantly different between the three populations (Figure 4b), with the fraction of CN variable base pairs per kilobase being greatest in *S. cerevisiae,* followed by *C. deuterogattii,* and then *S. pombe.* In contrast, the *S. cerevisiae* population has fewer SNPs per kilobase compared to *S. pombe* but more SNPs per kilobase compared to *C. deuterogattii* (Figure 4b).

How CN variants influence gene expression and phenotype in fungi is not well known. Examination of the contribution of CN variants to gene expression and phenotypic variation in *S. pombe* shows that partial aneuploidies (i.e., large CN variants) influence both local and global gene expression (Chikashige et al., 2007); in addition, CN variants are positively correlated with gene expression changes (*r_s_* = 0.71; *p* = 0.01; Spearman rank correlation; reported in Jeffares et al., 2017). Genome-wide association analyses of numerous phenotypes in *S. pombe* showed that structural variants accounted for 11% of phenotypic variation (CN variants accounted for 7% of that variation and rearrangements for 4%; Jeffares et al., 2017). The phenotypes significantly influenced by CN variants included growth rate, growth in various free amino acids (e.g., tryptophan, isoleucine), growth in the presence of various stressors (e.g., hydrogen peroxide, ultraviolet radiation, minimal media), and sugar utilization in winemaking (Jeffares et al., 2017). Although more studies are needed, these findings argue that CN variation may be a substantial contributor to the total genetic and phenotypic variation of fungal populations. Additionally, the variation in the correlation between CN and SNP variation across fungal populations (Figure 4) suggests that levels of SNP variation are not always a good proxy for levels of CN variation.

## Copy number variation and its impact on wine yeast adaptation in fermentation-related processes

During the wine making process, *S. cerevisiae* yeasts are barraged with numerous stressors such as high acidity, ethanol, osmolarity, sulfites, and low levels of oxygen and nutrient availability (Marsit and Dequin, 2015). Not surprisingly, *S. cerevisiae* strains isolated from wine making environments tend to be more robust to acid, copper, and sulfite stressors than yeasts isolated from beer and sake environments (Gallone et al., 2016). These biological differences are, at least partially, explained by variants, including CN variants, found at different frequencies or uniquely in wine yeasts. Below, we discuss what is known about the CN profile of genes from *S. cerevisiae* wine yeast strains associated with these stressors that may reflect diversity in stress tolerance or metabolic capacity and efficiency (Figure 5).

**Figure 5.**
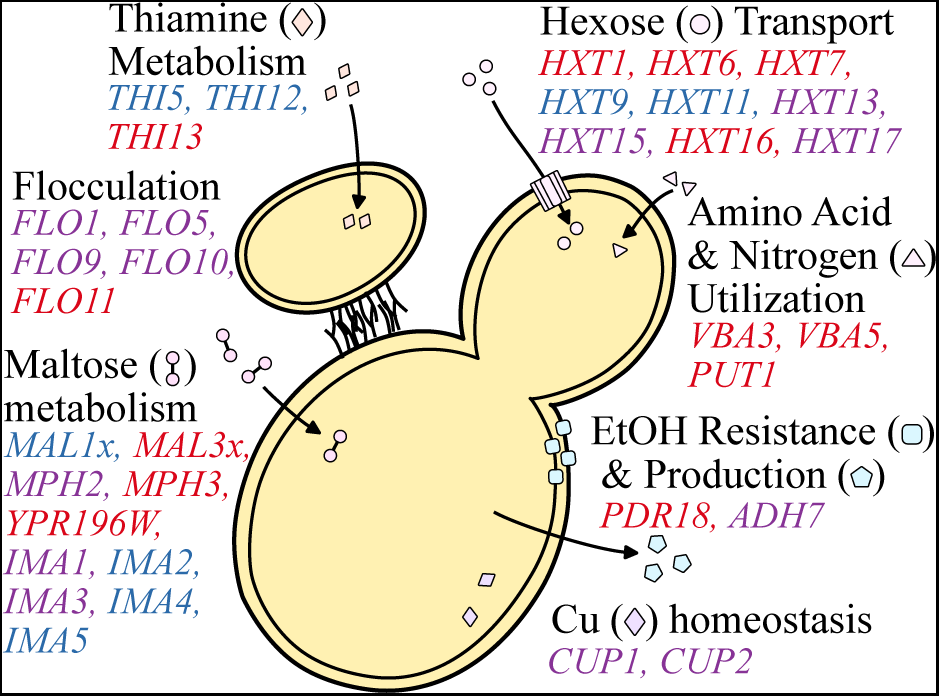
CN variable genes that affect functions important to wine making. Functional categories (e.g., Cu and Fe homeostasis, maltose metabolism, etc.) are shown in black font. Genes of interest are shown proximal to the category described and are colored blue, red, or purple to represent a gene observed to be primarily deleted, duplicated, or both across populations and studies investigating *S. cerevisiae* wine strains. Genes found to be both duplicated and deleted present an opportunity for oenologists to capitalize on standing genetic diversity to select for particular flavor profiles or yeast performance.

### CN variable genes related to stress

Many of the CN variable genes that have been identified among wine strains of *S. cerevisiae* (Gallone et al., 2016; Gonçalves et al., 2016; Ibáñez et al., 2014; Steenwyk and Rokas, 2017) are associated with fermentation processes (Table 1), which supports the hypothesis that CN variation plays a significant role in microbial domestication (Gibbons and Rinker, 2015). For example, *CUP1* is commonly duplicated among wine yeast strains, but not among yeasts in the closely related natural oak lineage (Almeida et al., 2015). Duplications in *CUP1* have been shown to confer copper resistance (Warringer et al., 2011) and their occurrence in wine yeast strains may have been driven by the human use of copper as a fungicide to combat powdery mildews in vineyards since the 1800’s (Almeida et al., 2015; Fay et al., 2004).

**Table 1.** Genes associated with fermentation-related processes that exhibit CN variation among *S. cerevisiae* wine strains

| Process<br>(organized<br>alphabetically) | Gene | Primarily<br>duplicated, deleted,<br>or both | Reference<br>(organized<br>alphabetically) |
| --- | --- | --- | --- |
| Amino acid &<br>nitrogen utilization | *VBA3, VBA5, PUT1* | Duplicated | Gallone *et al.* 2016;<br>Ibáñez *et al.* 2013 |
| Cu and Fe<br>homeostasis | *CUP1, CUP2* | Both | Almeida *et al.* 2015;<br>Fay *et al.* 2004;<br>Steenwyk & Rokas,<br>2017; Warringer *et al.* 2011 |
|  | *FIT2, FIT3, FRE3* | Duplicated | Gallone *et al.* 2016 |
| Ethanol resistance<br>and production | *PDR18* |  | Gallone *et al.* 2016 |
|  | *ADH7* | Both | Steenwyk & Rokas,<br>2017 |
| Flocculation | *FLO11* | Duplicated | Steenwyk & Rokas,<br>2017 |
|  | *FLO1, FLO5, FLO9,*<br>*FLO10* | Both | Gallone *et al.* 2016;<br>Steenwyk & Rokas,<br>2017 |
| Hexose transport | *HXT1, HXT4, HXT6,*<br>*HXT7, HXT16* | Duplicated | Gallone *et al.* 2016;<br>Steenwyk & Rokas,<br>2017 |
|  | *HXT9, HXT11* | Deleted | Gallone *et al.* 2016;<br>Steenwyk & Rokas,<br>2017 |
|  | *HXT13, HXT15,* | Both | Gallone *et al.* 2016; |
|  | *HXT17* |  | Steenwyk & Rokas, 2017 |
| Maltose metabolism | *MAL3x, MPH3, YPR196W* | Duplicated | Gallone *et al.* 2016; Gonçalves *et al.* 2016; Steenwyk & Rokas, 2017 |
|  | *MAL1x, IMA2, IMA4, IMA5* | Deleted | Gallone *et al.* 2016; Gonçalves *et al.* 2016; Steenwyk & Rokas, 2017 |
|  | *MPH2, IMA1, IMA3* | Both | Gallone *et al.* 2016; Steenwyk & Rokas, 2017 |
| Thiamine metabolism | *THI13* | Duplicated | Steenwyk & Rokas, 2017 |
|  | *THI5, THI12* | Deleted | Steenwyk & Rokas, 2017 |

Wine yeasts have also evolved strategies that favor survival in the wine fermentation environment, such as flocculation. This aggregation of yeast cells is associated with escape from hypoxic conditions, as it promotes floating and reaching the air-liquid interface where oxidative metabolism is possible (Fidalgo et al., 2006; Martínez et al., 1997). Flocculation is also favorable for oenologists as it facilitates yeast removal in post-processing (Soares, 2011) and is associated with the production of flavor enhancing ester-containing compounds (Pretorius, 2000). Flocculation is controlled by the *FLO* family of genes (Fidalgo et al., 2006; Govender et al., 2008). Examination of patterns of CN variation in *FLO* gene family members shows frequent duplications in *FLO11* as well as numerous duplications and deletions in *FLO1, FLO5, FLO9,* and *FLO10* (Gallone et al., 2016; Steenwyk and Rokas, 2017). Some of this variation may be adaptive. For example, partial duplications in the Serine/Threonine-rich hydrophobic region of *FLO11* are associated with the adaptive phenotype of floating to the air-liquid interface to access oxygen among “flor” or “sherry” yeasts (Fidalgo et al., 2006). Furthermore, the same partial duplications have also been observed in the more general wine population (Steenwyk and Rokas, 2017), suggesting that the benefits associated with this phenotype may not be unique to “flor” yeasts.

CN variation is also observed in genes related to stuck (incomplete) or sluggish (delayed) fermentations. Stuck fermentations are caused by a multitude of factors including nitrogen availability, nutrient transport, and decreased resistance to starvation (Salmon, 1989; Thomsson et al., 2005). Two genes associated with decrease resistance to starvation, *ADH7* and *AAD3,* are sometimes duplicated or deleted among wine yeast strains (Steenwyk and Rokas, 2017). Diverse CN profiles of *ADH7,* an alcohol dehydrogenase that reduces acetaldehyde to ethanol during glucose fermentation, and *AAD3,* an aryl-alcohol dehydrogenase whose null mutant displays greater starvation sensitivity (Walker et al., 2014), suggest variable degrees of starvation sensitivity and therefore fermentation performance. Additionally, wine yeasts are enriched for duplication in *PDR18* (Gallone et al., 2016), a transporter that aids in resistance to ethanol stress, one of the traits that differentiates wine from other industrial strains. Another gene associated with decreased resistance to starvation that also exhibits CN variation is *IMA1* (Steenwyk and Rokas, 2017), a major isomaltase with glucosidase activity (Teste et al., 2010).

### CN variable genes related to metabolism

Nutrient availability and acquisition is a major driving factor of wine fermentation outcome. Among the most important nutrients dictating the pace and success of wine fermentation is sugar availability (Marsit and Dequin, 2015). The most abundant fermentable hexose sugars in the wine environment include glucose and fructose (Marques et al., 2015), whose transport is largely carried out by genes from the hexose transporter (*HXT*) family (Boles and Hollenberg, 1997). A reproducible evolutionary outcome of yeasts exposed to glucose-limited environments, which are reflective of late wine fermentation, is duplication in hexose transporters, such as *HXT6* and *HXT7* (Brown et al., 1998; Dunham et al., 2002; Gresham et al., 2008, 2010), suggesting that changes in transporter CN are adaptive. Interestingly, the genes from the *HXT* gene family are highly CN variable among wine yeast strains (Dunn et al., 2012; Steenwyk and Rokas, 2017). For example, *HXT13, HXT15,* and *HXT17* exhibit CN variation among wine strains, *HXT1, HXT6, HXT7,* and *HXT16* are more commonly duplicated, and *HXT9* and *HXT11* are more commonly deleted (Gallone et al., 2016; Steenwyk and Rokas, 2017).

Similarly striking patterns of CN variation are observed for genes associated with maltose metabolism (Gallone et al., 2016; Gonçalves et al., 2016; Steenwyk and Rokas, 2017). The two *MAL* loci in the reference genome of *S. cerevisiae* S288C, *MAL1* and *MAL3,* that contain three genes which encode for a permease (*MALx1*), a maltase (*MALx2*), and a trans-activator (*MALx3*) (Michels et al., 1992; Naumov et al., 1994). The *MAL* loci are primarily associated with the metabolism of maltose (Michels et al., 1992) and therefore would be expected to be primarily deleted among wine yeasts as maltose is in relatively low abundance compared to other sugars during wine fermentation. As expected, the *MAL1* locus is deleted across many wine yeasts (Gallone et al., 2016; Gonçalves et al., 2016; Steenwyk and Rokas, 2017). In contrast, the *MAL3* locus is primarily duplicated among wine yeast strains (Gonçalves et al., 2016; Steenwyk and Rokas, 2017). Interestingly, part of the *MAL3* locus, *MAL32,* has been demonstrated to be important for growth on turanose, maltotriose, and sucrose (Brown et al., 2010), which are present in the wine environment, albeit in small quantities (M.Victoria and M. Carmen, 2013), suggesting potential function on secondary substrates or perhaps another function.

Equally important as sugar availability in determining fermentation outcome is nitrogen acquisition (Marsit and Dequin, 2015). Genes associated with amino acid and nitrogen utilization are commonly duplicated among wine yeast strains. Notable examples of such duplications are the amino acid permeases, *VBA3* and *VBA5* (Gallone et al., 2016), and *PUT1,* a gene that aids in the recycling or utilization of proline (Ibáñez et al., 2014).

CN variation is also observed in genes of the *THI* family, which are involved in thiamine, or vitamin B1, metabolism (Li et al., 2010), another important determinant of wine fermentation outcome. Several *THI* gene family members are CN variable; *THI5* and *THI12* are typically deleted, while *THI13* is commonly duplicated (Steenwyk and Rokas, 2017). Expression of *THI5* is commonly repressed or absent in wine strains, as it is associated with an undesirable rotten-egg smell and taste in wine (Bartra et al., 2010; Brion et al., 2014). Interestingly, *THI5* is deleted in greater than 90% of examined wine strains (Steenwyk and Rokas, 2017) but is duplicated in several other strains of *S. cerevisiae,* as well as in its sister species *S. paradoxus* and the hybrid species *S. pastorianus* (Wightman and Meacock, 2003).

## Conclusions and perspectives

An emerging body of work suggests that CN variation is an important, largely underappreciated, dimension of fungal genome biology and evolution (Farrer et al., 2013; Gallone et al., 2016; Gonçalves et al., 2016; Hartmann and Croll, 2017; Hu et al., 2011a; Steenwyk et al., 2016; Steenwyk and Rokas, 2017). Not surprisingly, numerous questions remain unresolved. For example, we have detailed numerous mechanisms that lead to the generation of CN variation but the relative contribution of each remains unclear. Additionally, both the genomic organization and genetic architecture of CN variants remain largely unknown. For example, are duplicated copies typically found in the same genomic neighborhood or are they dispersed? Similarly, what percentage of phenotypic differences among fungal strains is explained by CN variation?

The same can be said about the role of CN variation in yeast adaptation to the wine fermentation environment. Comparison of genome-wide patterns of CN variation among yeast populations responsible for the fermentation of different wines (e.g., white and red) would provide insight to how human activity has shaped the genome of yeasts associated with particular types of wine. Additionally, most sequenced wine strains originate from Italy, Australia, or France. Genome sequencing of yeasts from underrepresented regions (e.g., Africa and the Americas) may provide further insight to CN variable loci unique to each region and the global diversity of wine yeast genomes.

Another major set of questions are associated with examining the impact of CN variable loci at the different stages of wine fermentation. Insights on how CN variable loci modify gene expression, protein abundance and in turn fermentation behavior and end-product would be immensely valuable. A complementary, perhaps more straightforward, approach would be focused on examining the phenotypic impact of single-gene or gene family CN variants, such as the ones discussed in previous sections (e.g. genes belonging to the *ADH, HXT, MAL,* and *VBA* families; Table 1) on fermentation outcome. Such studies may provide an important bridge between scientist, oenologist, and wine-maker to enhance fermentation efficiency and consistency between batches or in the design of new wine flavor profiles.

Although this review focused solely on the contribution of *S. cerevisiae* CN variation, it is important to keep in mind that several other yeasts are also part of the wine fermentation environment. Members of many other wine yeast genera (e.g., *Hanseniaspora, Saccharomycodes,* and *Torulaspora*) are known to modify properties wine fermentation end product (Ciani and Maccarelli, 1998). Furthermore, recent sequencing projects have made several non-conventional wine yeast genomes publically available such as several *Hanseniaspora* species (Seixas et al., 2017; Sternes et al., 2016), *Starmerella bacillaris* (Lemos Junior et al., 2017), and *Lachancea lanzarotensis* (Sarilar et al., 2015). In-depth sequencing of populations from these yeast species and others associated with wine will provide insight to niche specialization within the wine environment as well as greatly enhance our understanding of CN variation and its role in the ecology and evolution of fungal populations.

## Abbreviations

BIR: break-induced recombination
CN: copy number
HR: homologous recombination
NHR: non-homologous repair
SNPs: single nucleotide polymorphisms

